# A theoretical approach to coupling the epithelial-mesenchymal transition (EMT) to extracellular matrix (ECM) stiffness via LOXL2

**DOI:** 10.1101/2021.02.04.429859

**Authors:** Youyuan Deng, Priyanka Chakraborty, Mohit Kumar Jolly, Herbert Levine

## Abstract

The epithelial-mesenchymal transition (EMT) plays a critical role in cancer progression, being responsible in many cases for the onset of the metastatic cascade and being integral in the ability of cells to resist drug treatment. Most studies of EMT focus on its induction via chemical signals such as TGF-β or Notch ligands, but it has become increasingly clear that biomechanical features of the microenvironment such as ECM (extracellular matrix) stiffness can be equally important. Here, we introduce a coupled feedback loop connecting stiffness to the EMT transcription factor ZEB1, which acts via increasing the secretion of LOXL2 that leads to increased cross-linking of collagen fibers in the ECM. This increased cross-linking can effectively increase ECM stiffness and increase ZEB1 levels, thus setting a positive feedback loop between ZEB1 and ECM stiffness. To investigate the impact of this non-cell-autonomous effect, we introduce a computational approach capable of connecting LOXL2 concentration to increased stiffness and thereby to higher ZEB1 levels. Our results indicate that this positive feedback loop, once activated, can effectively lock the cells in a mesenchymal state.

## Introduction

Metastasis remains the most lethal aspect of cancer progression. Various steps of the metastatic cascade have been intensely investigated from a biochemical signaling perspective. This has led to the concept of phenotypic plasticity – the ability of cells to reversibly switch their phenotypes in response to their environment – being identified as a hallmark of metastasis [1–3]. Recently, biophysical and biomechanical aspects of adaptability or plasticity have also begun to attract attention in the context of metastasis [4,5]. However, an integrated understanding of tumor cell plasticity and tissue mechanics remains largely elusive.

A key axis of phenotypic plasticity is the epithelial-mesenchymal transition (EMT) through which cells can alter their ability to migrate, invade, adhere to their neighbors and simultaneously evade attacks by drugs and the immune system [6]. Intracellular processes as well as cell-cell communication signaling networks underlying EMT dynamics have been mapped out extensively through the latest advances in high-throughput data acquisition such as RNA-seq, ATAC-seq, ChIP-seq and mass cytometry [7–11]. This data deluge has led to novel mechanism-based and data-based computational models to decode the underlying principles underlying the nonlinear dynamics of EMT [12–20]. Crucially, this analysis has revealed that EMT is not a binary process; instead, cells may occupy different positions in the high-dimensional space of molecular and/or morphological axes of EMT [21–23].

EMT is also influenced by tissue mechanics, and in turn, can influence the tissue mechanics [24]. Higher stiffness of extracellular matrix (ECM) can promote EMT and breast cancer invasion via the mechanosensitive EPHA2/LYN protein complex that facilitates nuclear localization of TWIST1, an EMT-inducing transcription factor (EMT-TF) [25,26]. In hepatocellular carcinoma, high stiffness can activate another EMT-TF SNAIL1, thus enabling invasion and metastasis [27]. Consistent observations are reported in oral squamous cell carcinoma and lung cancer cells [28,29]. On the other hand, EMT-TF ZEB1 can increase the protein levels of members of the lysyl oxidase (LOX) family of enzymes LOXL and LOXL2. Among other roles, LOXL2 crosslinks and stabilizes collagen to increase matrix stiffness, thus promoting lung cancer cell invasion and migration and eventually metastasis [30]. Consistently, LOXL2 has been associated with EMT and/or metastasis in colorectal cancer [31], gastric cancer [32], cervical cancer [33], and basal-like breast cancer [34]. Also, LOXL2 has been associated with enhanced chemotherapy resistance in triple negative breast cancer [35]. The pro-metastasis and drug-resistance role of LOXL2 has been attributed to increased levels of many EMT-TFs including SNAIL [34,36,37], which can activate ZEB1 [38]. At least in some cases, these effects appear to be directly due to the ECM remodeling aspects of LOXL2. Therefore, ZEB1 is involved in a non-cell-autonomous feedback loop where EMT and matrix stiffness can promote one another. However, the coupled dynamics of the EMT phenotype of a cell and the stiffness of the surrounding ECM has not yet been investigated.

In this work, we first use TCGA data to verify the general connection between high LOXL2, indicative of enhanced ECM stiffness, and ZEB1, indicative of enhanced metastatic capability and drug resistance. Then, we introduce a computational ECM model which allows for the evaluation of how secreted LOXL2 would influence matrix stiffness via its cross-linking capability. Finally, we use the aforementioned couplings to devise a circuit model which incorporates the ZEB1-LOXL2 mutual activation loop into existing systems biology treatments of EMT. Results from this augmented model indicate how the ECM feedback enhances the stability of more mesenchymal states in a cell density dependent manner and makes more difficult the reversion back to epithelial phenotypes

## II Methods

### LOXL2 correlation analysis

All the analysis was performed in R version 3.4.3 and correlations were calculated using cor() and “pearson” method in R. Three different EMT scoring methods – KS, MLR, 76GS were used to score samples separately in each dataset [39]. The TCGA datasets were downloaded from https://xenabrowser.net/datapages/. CCLE dataset was downloaded from https://portals.broadinstitute.org/ccle/data

### A mechanical model of collagen network

We model the extra-cellular matrices that are mainly composed of collagen as a 2D diluted, partially phantom, triangular lattice network (Figure 2(A)). The potential energy consists of terms that resists bond stretching and bending. Taking the spacing between nearest neighbor vertices as unity, then for the bond between *i*-th and *j*-th vertices,

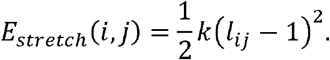

Two consecutive initially colinear bonds joined by a single vertex form a fiber segment, which has an elastic bending energy

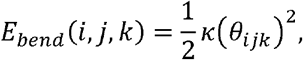

where *θ_ijk_* = 0 if bonds *i* − *j*, *j* − *k* are colinear. The total potential energy is *E_total_* = *E_stretch_* + *E_bend_*. Typically, the bond bending coefficient is taken to be much smaller than the bond stretching one, due to the small cross-sectional area of the individual fibers.

Each vertex acts as a hinge for different fibers to rotate freely. A non-phantom and non-diluted triangular lattice has a connection number of 6. That is, three fibers join at each vertex. Partial phantomness means that with probability *p_phan_*, a random one of the three fibers detach from the vertex. The lattice network is also diluted, i.e. each bond is formed only with a probability *p_bond_*. As such, the total average connectivity for a vertex is

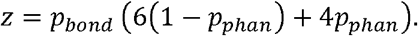

This network stiffens with increasing *z*, as demonstrated by the linear shear modulus depicted in Figure 2(B). To calculate the linear shear modulus, we apply a small strain *γ* with Lee-Edwards periodic boundary condition (i.e. by horizontally shifting neighboring periodic images above and below the computational box), and thereafter relax the network by minimizing *E_total_* = *E_stretch_* + *E_bend_*. The linear shear modulus is then 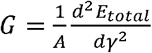, where A is the area per period. The same formula for *G*(*γ*) is also applicable in the nonlinear regime of large strains.

### Modeling local crosslink and stiffness variation in extra-cellular matrices

LOXL2 promotes crosslinks between fibers. We take this into account by assuming a direct correspondence between a local *p_phan_* and the local LOXL2 concentration. This decrease in phantomization mimicking the increased cross-linking increases the connectivity *z* and hence the stiffness. To measure the local stiffness in this lattice just described, we perform bead experiments *in silico* as in [40]. To measure the stiffness at some location, we embed a bead therein and apply external harmonic “optical traps” along four orthogonal directions, ±*x*, ±*y*. The extra potential energy due to the trap is

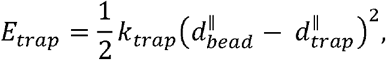

where 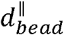 and 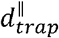 are the position of the bead and trap bottom respectively, measured with respect to the initial location of the bead along x or y axis whichever is parallel to the trap gradient. The total potential energy *E_total_* = *E_stretch_* +*E_bend_* + *E_trep_* is then minimized with respect to the bead and vertex degrees of freedom. The local compliance (inversely related to stiffness) is defined as

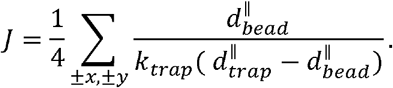

Such a measurement can be repeated at different locations on the same network, one bead at a time. Fixed boundary conditions are applied; using a periodic boundary as in the bulk shearing would result in the whole lattice translating with no energy penalty. To avoid artifacts due to edge effects, the measurement is only performed on a slightly smaller inner region of the whole lattice.

### Modeling positive mutual feedback between cells and ECM

We build on the previously established set of ordinary differential equations for the core EMT biochemical network to include the new feedback loop discussed in this text (Figure 4(A)). We solve for the steady state under different levels of external signaling and draw the bifurcation diagram using the software Matcont (https://sourceforge.net/projects/matcont/) with a custom Matlab script.

## III Results

### LOXL2 correlates positively with EMT markers and a more mesenchymal signature

First, we investigated the correlation between the expression levels of LOXL2 and that of key EMT markers for tumor samples, using various TCGA datasets (BRCA, COAD, COADREAD, OV), and for cancer cell lines using the Cancer Cell Line Encyclopedia (CCLE) dataset. LOXL2 consistently correlated positively with ZEB1 but negatively with CDH1 (a standard epithelial marker), across all datasets (Figure 1).

**Figure 1:**
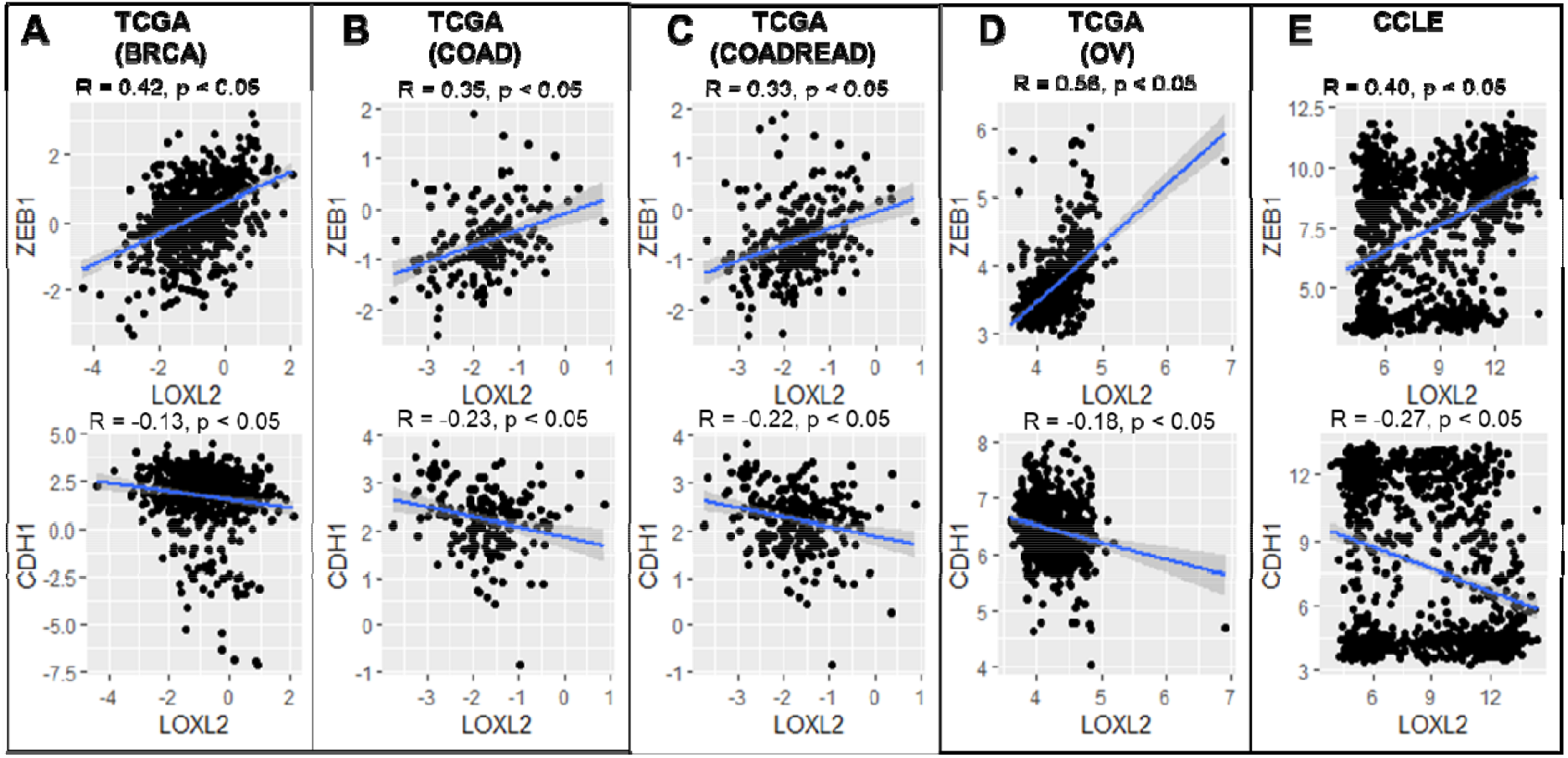
Correlation between EMT markers and LOXL2 gene expression values. Pearson’s correlation values (R, p) are bee included; the regression line has been highlighted in blue.

**Figure 2:**
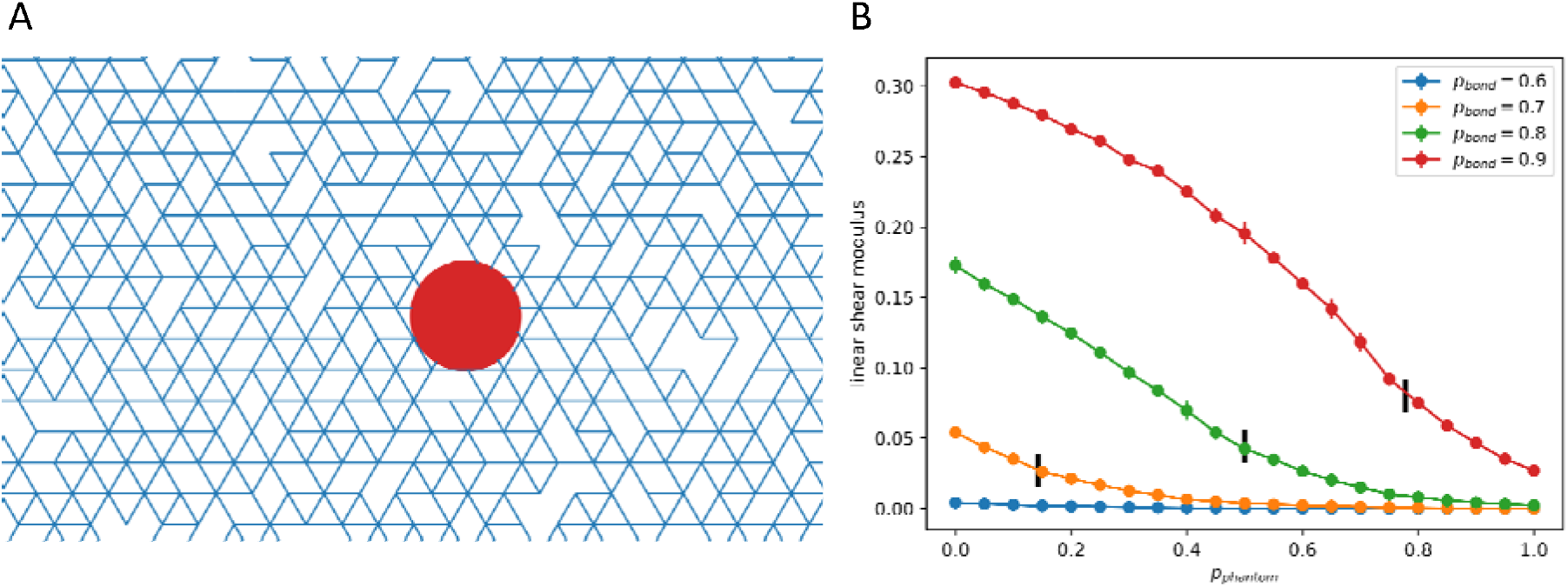
(A) A local region of the diluted triangular lattice. The red circle represents an embedded bead used for measuring local stiffness. (B) Linear shear modulus for different p_phan_ and p_bond_, black bars represent the point where average connectivity equals to 4.

Next, we calculated the EMT scores for these cell lines and tumor samples using three different EMT scoring methods (76GS, KS, MLR) and checked their correlation with LOXL2. The 76 GS method has no specific pre-defined range of values and a higher 76GS score associates with an epithelial phenotype. Unlike the 76 GS method, the MLR and KS methods have predefined scales for EMT scores. MLR and KS score EMT on a range of [0, 2] and [−1, 1] respectively, with higher scores indicating a mesenchymal phenotype. The 76GS score showed a negative correlation with LOXL2 and as expected, the other two scores (KS, MLR) showed a positive correlation with LOXL2 across all tumor datasets (Figure S1). This analysis signifies that the LOXL2 gene expression levels are associated with a more mesenchymal phenotype in tumor datasets. However, these patterns were absent in the CCLE dataset.

### Cross-linking effect of LOXL2 stiffens the collagen network

Given this correlation, we were motivated to develop a modeling framework that could account for this effect given the known interactions between these molecular players. As already mentioned, it is well-known that LOXL2 can increase the stiffness of extracellular matrix by cross-linking collagen. In recent years, there has been considerable effort devoted to formulating computational models of fibrous matrices typical of what would be expected in the stromal regions surrounding tumors. One popular approach is to consider a diluted lattice representation of the mechanics (triangular for a 2D system, face-centered-cubic in 3D) with both stretching and bending energies governing deformations of each link [41,42]. An example of one such diluted triangular lattice in 2D, is shown in Figure 2(A). As discussed at length in the literature, the mechanics of this model can be highly non-linear, exhibiting strong shear-stiffening [41]. Mackintosh and co-workers have extended this basic approach to take into account variations in cross-linking [43,44]. Specifically, they introduced the idea of phantom crossings, sites at which some fibers (i.e., the continuing links) that pass through that site do not interact with other fibers.

For all the aforementioned lattice mechanical models, there exists a special critical value of the average connectivity *z* (number of bonds connected) of each lattice point, the so-called isostatic point *z_IP_*, whose value equals two times the dimensionality. ECM that is under normal physiological condition is considered to be below the isostatic point. For example, in the 2D models by Mackintosh and co-workers, there is always full phantomization. i.e. every site is phantomized such that the connectivity is always below 4 = 2 × 2. This then gives rise to the aforementioned strain-stiffening and explains similar findings from experiments on reconstituted collagen I networks [43,44].

We adopt the idea of phantomization but make it a tunable, in order to capture the upregulated cross-linking in cancerous tissues. For simplicity, we do this using the 2D triangular lattice framework (see Methods) as the basic physics of the stress response is independent of spatial dimension [44]. If the probability of having a node with this assumed “phantom-like” behavior is *p_phan_* and the probability of a link being present is *p_bond_*, the average connectivity becomes

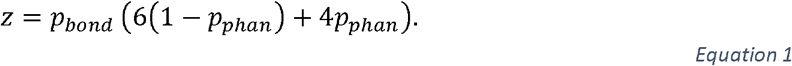

As already mentioned, when *z* is below the isostatic point *z_IP_* = 4, the network stiffens only with sufficiently large strain by entering the nonlinear regime; this is the physiological strain-stiffening. Above this point, the network exhibits large stiffness even in the small-strain linear regime. (Reference [44], Figure 1 therein, presents an illustrative diagram.) Either by stress-stiffening or by increasing above this point, the stiffness becomes dominated by the (large) stretching stiffness and roughly independent of the (small) bending stiffness. In this work we focus on the increase in *z* by decreasing *p_phan_* (via LOXL2) enabling the linear regime to attain high stiffness, as what is likely in cancerous pathology. Note also that within the correct range of parameters, *p_bond_* can be varied to account for varying fiber density.

Under this framework, we model the molecular effect of LOXL2 as causing a decrease in the value of *p_phan_*. Figure 2(B) shows a set of response curves for the stiffness as we vary this parameter for different values of *p_bond_*. As can be seen, the stiffness exhibits a significant rise once we have exceeded the critical value *z_IP_* by sufficiently lowering *p_phan_*. This means that as LOXL2 concentration is increased past a threshold value, it will give rise to an increasingly stiff ECM. As the threshold clearly decreases with *p_bond_*, it will be easier to create stiff ECM at higher values of fiber density.

### Effects of spatial distribution of LOXL2

To get some idea of what might happen in practice for a localized tumor, we imagine that cells in some spherical region of radius *R* form a tumor of a constant cell density *ρ* and are emitting LOXL2 at a constant rate *α*. These molecules then diffuse with diffusion constant *D* and also decay at rate *λ*; in steady state, this creates an exponentially decaying concentration once we leave the tumor mass itself, namely

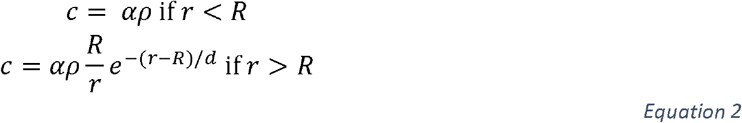

where the decay rate is 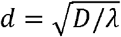. By assuming that this concentration modulates *p_phan_* via

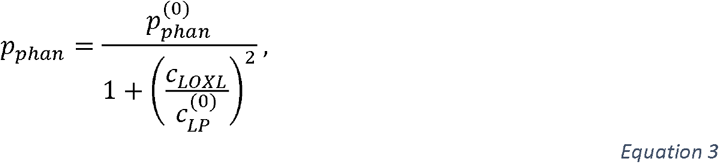

we can obtain a map of the factor by which the local compliance is decreased, corresponding to stiffness being increased, due to the cross-linking effect (See Methods). One such map for nominal parameter values is depicted in Figure 3. It shows how ECM outside of the tumor boundary is also stiffened, and demonstrates a potential non-cell-autonomous biomechanical effect on cells neighboring the bulk tumor. It would be useful in the future to measure this behavior directly by monitoring both concentration and compliance maps for an *in vitro* tumor spheroid.

**Figure 3:**
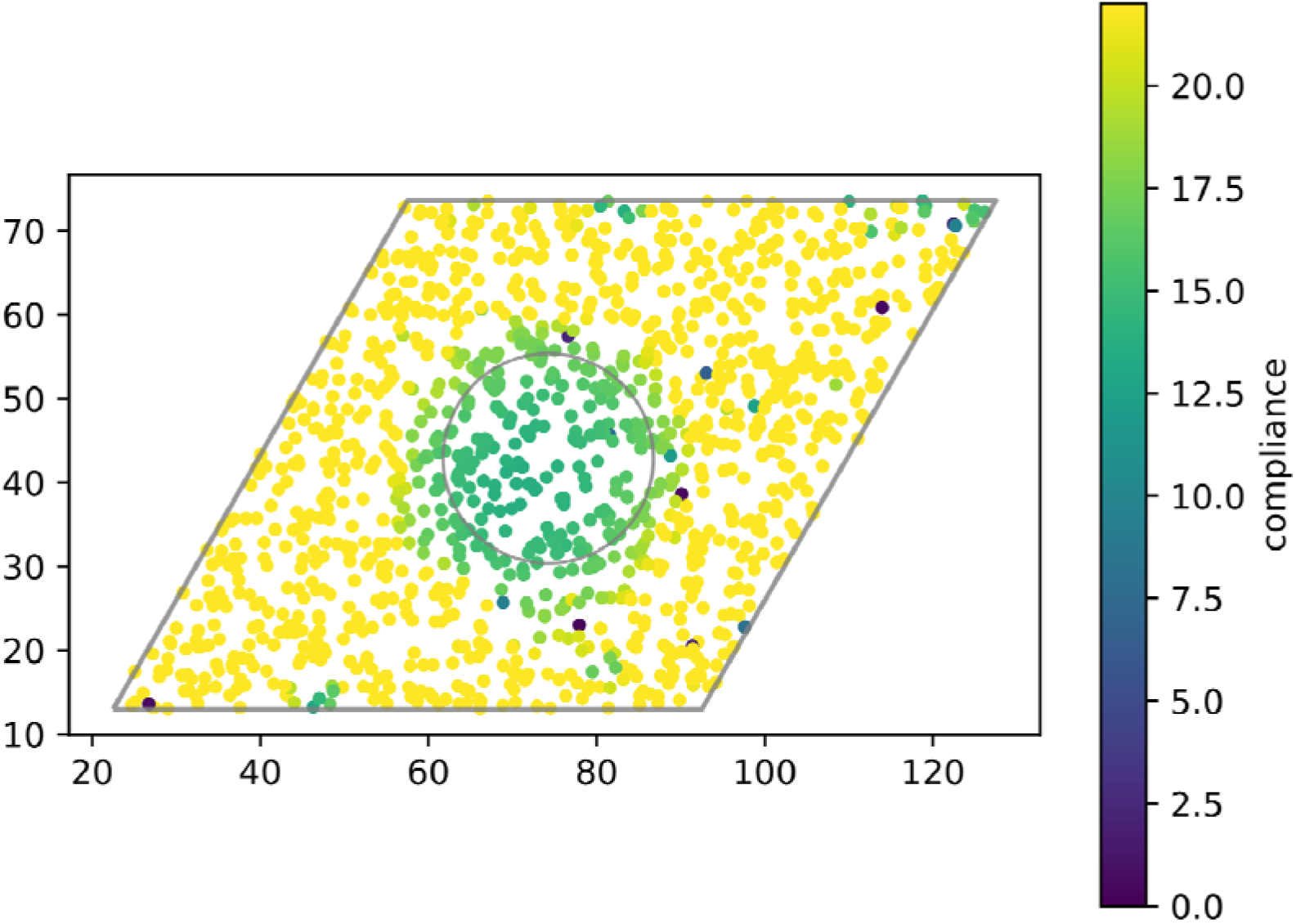
Local compliance measurement predicted by the triangular lattice network model. Cells inside the inner circles secrete LOXL2, which is then subject to diffusion and degradation, and modifies local collagen network connectivity. The outer grey parallelogram signifies the measured region, which is some distance away from the true computational boundary to avoid edge effects. The network is not rectangular because it follows the base vectors of the triangular unit cell.

**Figure 4:**
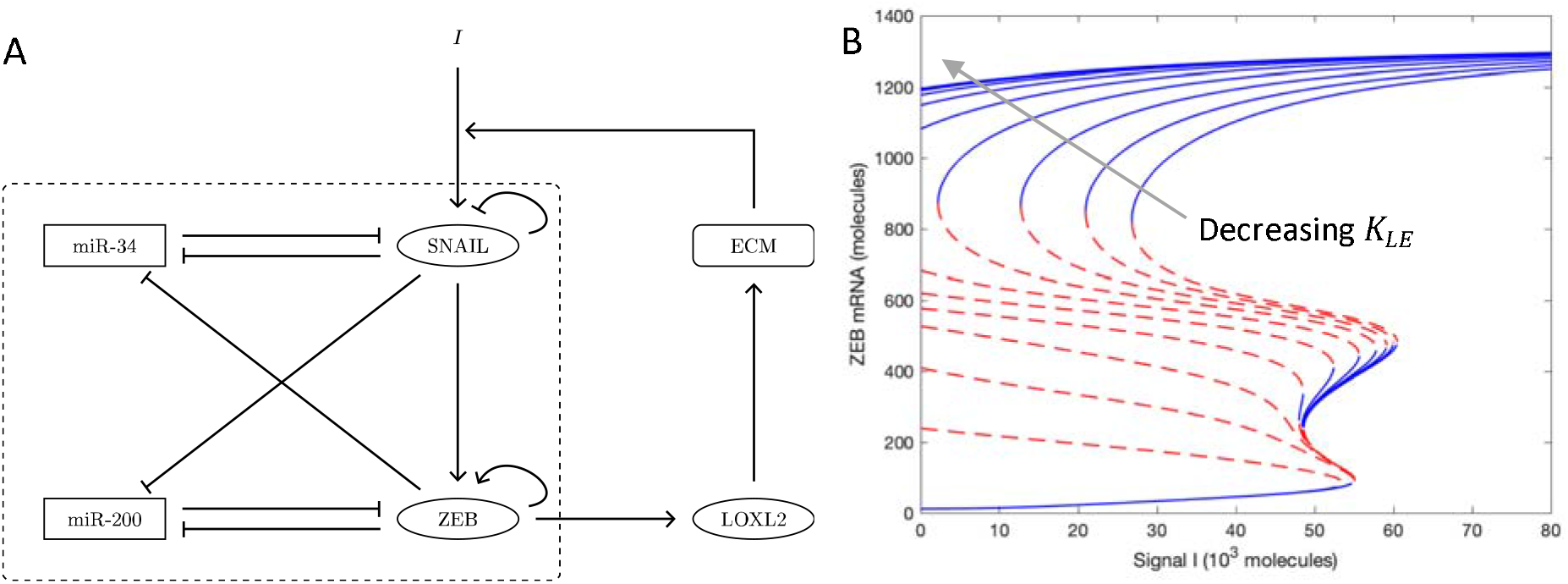
(A) Biochemical reaction network of core EMT circuit (within dashed box) plus the LOXL2- and ECM-mediated feedback loop including LOXL2. (B) Bifurcation diagrams of EMT core circuit coupled to the LOXL2 feedback loop. Different curves following the arrow correspond to different thresholds K_LE_.

### LOXL2-mediated feedback loop drives EMT

The ECM stiffness can couple directly to EMT phenotype via a variety of signaling pathways [25,28,45]. Elevated LOXL2 level increases ECM crosslinking and the increased mechanical stiffness is then input to SNAIL, as indicated by the mechanosensing studies mentioned above [25,26]. Thus, we can expect that as the matrix stiffens cells will assume more of a mesenchymal phenotype, thereby increasing the production levels of the ZEB1 transcription factor. Further, [30] shows that increasing ZEB1 will upregulate LOXL2. There is then a positive feedback loop which is expected to stabilize mesenchymal phenotypes. To demonstrate this effect, we implemented this feedback loop in our previously validated model of the core circuit governing EMT that consists of two interconnected, double-negative feedback loops and exhibits epithelial, hybrid, mesenchymal states [46].

We formulate the regulation from ZEB to LOXL2 using Michaelis–Menten kinetics:

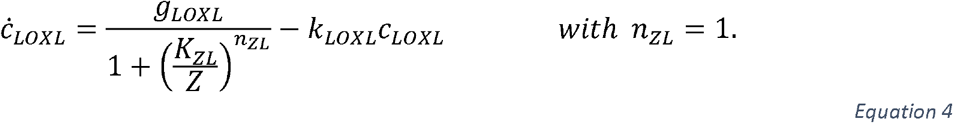

We neglect any delays in the ECM modification by LOXL2, and instead make *p_phan_* directly depend on LOXL2 levels. Since the stiffness is directly dependent upon *p_phan_*, we effectively input a mechanosensing signal to SNAIL:

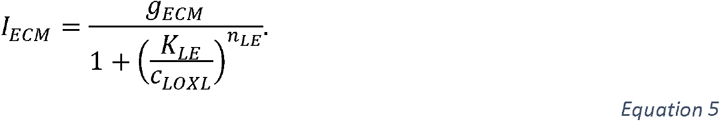

Although we can calculate the curve of ECM stiffness vs *p_phan_* (Figure 2(B)), the precise quantitative form of mechanosensing is unknown. This choice of a Hill function captures the fact that, for some intermediate *p_bond_*, *p_phan_* must decrease beyond the isostatic point before the mechanical lattice network stiffens. Qualitatively, Equation 5 combines Equation 3, Figure 2(B), and the unknown quantitative form of mechanosensing. The reaction network is displayed in Figure 4(A). Readers looking for the detailed parametrization are redirected to the SI.

The threshold constant *K_LE_* in the Hill function (Equation 5) captures the strength of mechanosensing feedback. Note that the absolute concentration of LOXL2 is irrelevant; only the ratio 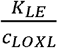 matters. As shown in Figure 2(B), ECM of higher density stiffens more easily; it starts to have high stiffness with lower concentration of LOXL2. This is equivalent to a smaller threshold *K_LE_*. Interpreted differently, *K_LE_* can be viewed as the “measuring stick” for *c_LOXL_*. A smaller *K_LE_* amounts to a larger 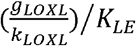 using this “measuring stick”. Since the ordinary differential equations are valid both for a single cell or for a group of cell of similar expression state, increasing LOXL2 could be attributed to higher secretion rates or higher cell density of the secreting cells. Specifically, if we assume that all nearby cells are in the same state, then the overall LOXL2 level scales with the cell density multiplied by the single cell secretion.

Shown in Figure 4(B) is a family of steady-state bifurcation curves of ZEB1 vs external signal *I*, going from right to left with decreasing *K_LE_*. A few features are worth noting. First, the epithelial states with very low ZEB values are not altered in any significant manner; except for the very extreme case where the mechanosensing signal is always at maximum regardless of LOXL2 concentration, only the mesenchymal state survives (whereby *K_LE_* = 0, see SI Figure S3). Next, the hybrid E/M states become increasingly rare as the feedback is ramped up. This is because having an intermediate value of ZEB is untenable if it can take part in the new positive feedback loop we have identified. Finally, there is a large hysteretic effect whereby mesenchymal states can remain stable even as the inducing signal is turned off, as M cells bootstrap their own existence by increasing the ECM stiffness in their immediate vicinity. This type of dynamical effect has previously been suggested as a consequence of TGF-β secretion by mesenchymal cells [47,48] as well as a possible “epigenetic locking” of cells in a mesenchymal state [49,50], but the results here are more dramatic because of the very strong nonlinearity of the stress response.

## IV Discussion

Cancer metastasis is one of the most vicious traits of cancer. The ability to metastisize is related to the plasticity of cancer cells, of which the epithelial-mesenchymal transition (EMT) is a key axis. Recently, it has been shown that in several cancer types, there can exist a possible positive feedback loop along the SNAIL1-ZEB1-LOXL2 axis [30–38]. Elevated levels of all three molecules are correlated with cancer malignancy [51–55]. Among them, LOXL2 is downstream to ZEB1 and has a known extra-cellular effect. It promotes crosslinking between collagen fibers, a major component of the extra-cellular matrix (ECM), thus stiffening and stabilizing the ECM. Increased stiffness can be fed back to SNAIL1 (i.e. increased stiffness can upregulate SNAIL levels) and further drives EMT. In this article, we have examined the bioinformatics-based, biochemical, and mechanical aspects of this feedback loop. We do note that there can be additional intracellular effects of LOXL2 [36]. These are not taken into account in our circuit as they do not represent a significant change to the original core circuit with its assumed self-activation positive feedback loop of ZEB.

Studying gene expression patterns from TCGA and CCLE database revealed a positive correlation between LOXL2 and ZEB1, as well as a negative correlation between LOXL2 and E-Cadherin. These findings are consistent with the notion that LOXL2 drives EMT and with the possibility that this driving is through the mechanical effect of increased collagen cross-linking. To demonstrate this possibility, we adapted the triangular lattice network model typically used to study the mechanical properties of collagen networks (including local stiffness). Specifically, we associated bond dilution with collagen density, and vertex phantomization with collagen crosslinks. Under small strain in the linear regime, this network stiffens significantly only when the connectivity is increased past two times the dimensionality, in our 2D case, four. A simplified biophysical setup is simulated, where a geometrically localized cluster of cancer cells secretes LOXL2 that is then subject to diffusion and degradation. In the steady state, increased LOXL2 and stiffness can “seep” outside the range of the cluster. From the biological perspective, this is a non-cell-autonomous effect.

To have a more quantitative picture, we incorporated this interaction into the core EMT regulatory network previously developed [46]. The input of the mechanosensing signal is assumed to take the form of a high order Hill activation function. Different threshold constants in the Hill function thereby correspond to how much the secreted LOXL2 can stiffen the ECM. Under circumstances that can correspond to a very low collagen density, increased cross-linking by LOXL2 barely stiffens the network, the corresponding Hill function always stays below threshold, and the EMT bifurcation diagram is hardly affected. Under circumstances on the opposite extreme, the bifurcation is strongly shifted such that the purely mesenchymal states stabilize and dominate.

Another factor that can affect the previous result is the population structure of the cells. The fact that stiffness interacts with EMT provides a mechanism for cells to affect each other; a population of mesenchymal cells can push a nearby cell to also undergo EMT. Although it is in principle possible that a single cell could secrete enough LOX2 to alter its own microenvironment, it seems more likely that this feedback sets in only in the presence of a sufficiently numerous population of cells. This implies that just as in the case of Notch signaling, the single-cell picture must give way to a more comprehensive tissue-scale analysis of EMT phenotypes and their spatial correlation [56,57].

Our modeling results offer an explanation of the experimentally observed correlations between the EMT-related transcriptional factors and extracellular enzymes. It also should inspire future studies integrating mechanics and cancer-related biology both experimentally and theoretically. Especially, we think that more accurate experiments mapping local matrix stiffness and ECM-modifying enzyme concentrations can test our model and can also open the door to a better understanding of the detailed roles of mechanical properties in cancer development. Nonetheless, the current approach is not without limitations. For example, we focus on the cross-linking by LOXL2 while keeping the collagen deposition density as an external factor. However, there is evidence that cancer cells can also promote collagen deposition [30]. The diluted, partially phantom lattice network described in this work can encompass this additional effect via varying the bond dilution factor. We also could include the aforementioned intracellular role of LOXL2 by extending our baseline EMT circuit. These extensions will become useful once more quantitative data becomes available connecting the various pieces of this novel feedback mechanism.

## Funding

MKJ was supported by InfoSys Foundation, Bangalore, and by Ramanujan Fellowship awarded by Science and Engineering Research Board (SERB), Department of Science and Technology (DST), Government of India (SB/S2/RJN-049/2018). HL and YD are supported by by NSF (PHY-1935762 and NSF (PHY-2019745).

## Notes

### Competing Interest Statement

The authors have declared no competing interest.

## References

1. Mooney, S.; Jolly, M.; Levine, H.; Kulkarni, P. Phenotypic plasticity in prostate cancer: Role of intrinsically disordered proteins. Asian J. Androl. 2016, 18, 704–710, doi:10.4103/1008-682X.183570.

2. Jolly, M.K.; Celia-Terrassa, T. Dynamics of Phenotypic Heterogeneity Associated with EMT and Stemness during Cancer Progression. J Clin Med 2019, 8, 1542, doi:10.3390/jcm8101542.

3. Celià-Terrassa, T.; Kang, Y. Distinctive properties of metastasis-initiating cells. Genes Dev. 2016, 30, 892–908, doi:10.1101/gad.277681.116.892.

4. Gensbittel, V.; Krater, M.; Harlepp, S.; Busnelli, I.; Guck, J.; Goetz, J.G. Mechanical Adaptability of Tumor Cells in Metastasis. Dev. Cell 2020, in press, doi:10.1016/j.devcel.2020.10.011.

5. Follain, G.; Herrmann, D.; Harlepp, S.; Hyenne, V.; Osmani, N.; Warren, S.C.; Timpson, P.; Goetz, J.G. Fluids and their mechanics in tumour transit: shaping metastasis. Nat. Rev. Cancer 2020, 20, 107–124, doi:10.1038/s41568-019-0221-x.

6. Jia, D.; Li, X.; Bocci, F.; Tripathi, S.; Deng, Y.; Jolly, M.K.; Onuchic, J.N.; Levine, H. Quantifying cancer epithelial-mesenchymal plasticity and its association with stemness and immune response. arXiv 2019.

7. Johnson, K.S.; Hussein, S.; Xong, S.; Chakraborty, P.; Jolly, M.K.; Toneff, M.J.; Lin, Y.C.; Taube, J.H. Gene expression and chromatin accessibility during progressive EMT and MET linked to dynamic CTCF engagement. bioRxiv 2020, 089110, doi:10.1101/2020.05.11.089110.

8. Malouf, G.G.; Taube, J.H.; Lu, Y.; Roysarkar, T.; Panjarian, S.; Estecio, M.R.H.; Jelinek, J.; Yamazaki, J.; Raynal, N.J.M.; Long, H.; et al. Architecture of epigenetic reprogramming following Twist1-mediated epithelial-mesenchymal transition. Genome Biol. 2013, 14, R144, doi:10.1186/gb-2013-14-12-r144.

9. Cook, D.P.; Vanderhyden, B.C. Context specificity of the EMT transcriptional response. Nat. Commun. 2020, 11, 2142, doi:10.1038/s41467-020-16066-2.

10. Karacosta, L.G.; Anchang, B.; Ignatiadis, N.; Kimmey, S.C.; Benson, J.A.; Shrager, J.B.; Tibshirani, R.; Bendall, S.C.; Plevritis, S.K. Mapping Lung Cancer Epithelial-Mesenchymal Transition States and Trajectories with Single-Cell Resolution. Nat. Commun. 2019, 10, 5587, doi:10.1101/570341.

11. Pastushenko, I.; Brisebarre, A.; Sifrim, A.; Fioramonti, M.; Revenco, T.; Boumahdi, S.; Van Keymeulen, A.; Brown, D.; Moers, V.; Lemaire, S.; et al. Identification of the tumour transition states occurring during EMT. Nature 2018, 556, 463–468, doi:10.1038/s41586-018-0040-3.

12. Hari, K.; Sabuwala, B.; Subramani, B.V.; Porta, C. La; Zapperi, S.; Font-Clos, F.; Jolly, M.K. Identifying inhibitors of epithelial-mesenchymal plasticity using a network topology based approach. npj Syst. Biol. Appl. 2020, 6, 15, doi:10.1038/s41540-020-0132-1.

13. Sha, Y.; Wang, S.; Zhou, P.; Nie, Q. Inference and multiscale model of epithelial-to-mesenchymal transition via single-cell transcriptomic data. Nucleic Acids Res. 2020, 48, 9505–9520, doi:10.1093/nar/gkaa725.

14. Devaraj, V.; Bose, B. Morphological State Transition Dynamics in EGF-Induced Epithelial to Mesenchymal Transition. J. Clin. Med. 2019, 8, 911, doi:10.3390/jcm8070911.

15. Mandal, M.; Ghosh, B.; Anura, A.; Mitra, P.; Pathak, T.; Chatterjee, J. Modeling continuum of epithelial mesenchymal transition plasticity. Integr. Biol. 2016, 8, 167–176, doi:10.1039/C5IB00219B.

16. Bocci, F.; Jolly, M.K.; Onuchic, J.N. A biophysical model uncovers the size distribution of migrating cell clusters across cancer types. Cancer Res. 2019, 79, 5527–5535, doi:10.1158/0008-5472.CAN-19-1726.

17. Font-Clos, F.; Zapperi, S.; Porta, C.A.M. La Topography of epithelial–mesenchymal plasticity. Proc. Natl. Acad. Sci. 2018, 115, 5902–5907, doi:10.1073/pnas.1722609115.

18. Zadran, S.; Arumugam, R.; Herschman, H.; Phelps, M.E.; Levine, R.D. Surprisal analysis characterizes the free energy time course of cancer cells undergoing epithelial-to-mesenchymal transition. Proc. Natl. Acad. Sci. U. S. A. 2014, 111, 13235–40, doi:10.1073/pnas.1414714111.

19. Xu, S.; Ware, K.; Ding, Y.; Kim, S.; Sheth, M.; Rao, S.; Chan, W.; Armstrong, A.; Eward, W.; Jolly, M.; et al. An Integrative Systems Biology and Experimental Approach Identifies Convergence of Epithelial Plasticity, Metabolism, and Autophagy to Promote Chemoresistance. J. Clin. Med. 2019, 8, 205, doi:10.3390/jcm8020205.

20. McFaline-Figueroa, J.L.; Hill, A.J.; Qiu, X.; Jackson, D.; Shendure, J.; Trapnell, C. A pooled single-cell genetic screen identifies regulatory checkpoints in the continuum of the epithelial-to-mesenchymal transition. Nat. Genet. 2019, 51, 1389–1398, doi:10.1038/s41588-019-0489-5.

21. Jolly, M.K.; Somarelli, J.A.; Sheth, M.; Biddle, A.; Tripathi, S.C.; Armstrong, A.J.; Hanash, S.M.; Bapat, S.A.; Rangarajan, A.; Levine, H. Hybrid epithelial/mesenchymal phenotypes promote metastasis and therapy resistance across carcinomas. Pharmacol. Ther. 2019, 194, 161–184, doi:10.1016/j.pharmthera.2018.09.007.

22. Goetz, H.; Melendez-Alvarez, J.R.; Chen, L.; Tian, X.-J. A plausible accelerating function of intermediate states in cancer metastasis. PLOS Comput. Biol. 2020, 16, e1007682, doi:10.1371/journal.pcbi.1007682.

23. Cheung, K.J.; Ewald, A.J. Illuminating breast cancer invasion: diverse roles for cell-cell interactions. Curr. Opin. Cell Biol. 2014, 30, 99–111, doi:10.1016/j.ceb.2014.07.003.

24. Coban, B.; Bergonzini, C.; Zweemer, A.J.M.; Danen, E.H.J. Metastasis: crosstalk between tissue mechanics and tumour cell plasticity. Br. J. Cancer 2020, in press.

25. Fattet, L.; Jung, H.-Y.; Matsumoto, M.W.; Aubol, B.E.; Kumar, A.; Adams, J.A.; Chen, A.C.; Sah, R.L.; Engler, A.J.; Pasquale, E.B.; et al. Matrix Rigidity Controls Epithelial-Mesenchymal Plasticity and Tumor Metastasis via a Mechanoresponsive EPHA2/LYN Complex. Dev. Cell 2020, 54, 302–316.e7, doi:10.1016/j.devcel.2020.05.031.

26. Wei, S.C.; Fattet, L.; Tsai, J.H.; Guo, Y.; Pai, V.H.; Majeski, H.E.; Chen, A.C.; Sah, R.L.; Taylor, S.S.; Engler, A.J.; et al. Matrix stiffness drives epithelial–mesenchymal transition and tumour metastasis through a TWIST1–G3BP2 mechanotransduction pathway. Nat. Cell Biol. 2015, 17, 678–688, doi:10.1038/ncb3157.

27. Dong, Y.; Zheng, Q.; Wang, Z.; Lin, X.; You, Y.; Wu, S.; Wang, Y.; Hu, C.; Xie, X.; Chen, J.; et al. Higher matrix stiffness as an independent initiator triggers epithelial-mesenchymal transition and facilitates HCC metastasis. J. Hematol. Oncol. 2019, 12, 112, doi:10.1186/s13045-019-0795-5.

28. Matte, B.F.; Kumar, A.; Placone, J.K.; Zanella, V.G.V.G.; Martins, M.D.; Engler, A.J.; Lamers, M.L. Matrix stiffness mechanically conditions EMT and migratory behavior of oral squamous cell carcinoma. J. Cell Sci. 2019, 132, jcs224360, doi:10.1242/jcs.224360.

29. Kim, D.; You, E.; Jeong, J.; Ko, P.; Kim, J.W.; Rhee, S. DDR2 controls the epithelial-mesenchymal-transition-related gene expression via c-Myb acetylation upon matrix stiffening. Sci. Rep. 2017, 7, 6847, doi:10.1038/s41598-017-07126-7.

30. Peng, D.H.; Ungewiss, C.; Tong, P.; Byers, L.A.; Wang, J.; Canales, J.R.; Villalobos, P.A.; Uraoka, N.; Mino, B.; Behrens, C.; et al. ZEB1 induces LOXL2-mediated collagen stabilization and deposition in the extracellular matrix to drive lung cancer invasion and metastasis. Oncogene 2017, 36, 1925–1938, doi:10.1038/onc.2016.358.

31. Park, P.G.; Jo, S.J.; Kim, M.J.; Kim, H.J.; Lee, J.H.; Park, C.K.; Kim, H.; Lee, K.Y.; Kim, H.; Park, J.H.; et al. Role of LOXL2 in the epithelial-mesenchymal transition and colorectal cancer metastasis. Oncotarget 2017, 8, 80325–80335, doi:10.18632/oncotarget.18170.

32. Peng, L.; Ran, Y.-L.; Hu, H.; Yu, L.; Liu, Q.; Zhou, Z.; Sun, Y.-M.; Sun, L.-C.; Pan, J.; Sun, L.-X.; et al. Secreted LOXL2 is a novel therapeutic target that promotes gastric cancer metastasis via the Src/FAK pathway. Carcinogenesis 2009, 30, 1660–1669, doi:10.1093/carcin/bgp178.

33. Tian, J.; Sun, H.-X.; Li, Y.-C.; Jiang, L.; Zhang, S.-L.; Hao, Q. LOXL 2 Promotes The Epithelial–Mesenchymal Transition And Malignant Progression Of Cervical Cancer. Onco. Targets. Ther. 2019, 12, 8947–8954, doi:10.2147/OTT.S217794.

34. Moreno-Bueno, G.; Salvador, F.; Martín, A.; Floristán, A.; Cuevas, E.P.; Santos, V.; Montes, A.; Morales, S.; Castilla, M.A.; Rojo-Sebastián, A.; et al. Lysyl oxidase-like 2 (LOXL2), a new regulator of cell polarity required for metastatic dissemination of basal-like breast carcinomas. EMBO Mol. Med. 2011, 3, 528–544, doi:10.1002/emmm.201100156.

35. Saatci, O.; Kaymak, A.; Raza, U.; Ersan, P.G.; Akbulut, O.; Banister, C.E.; Sikirzhytski, V.; Tokat, U.M.; Aykut, G.; Ansari, S.A.; et al. Targeting lysyl oxidase (LOX) overcomes chemotherapy resistance in triple negative breast cancer. Nat. Commun. 2020, 11, 2416, doi:10.1038/s41467-020-16199-4.

36. Salvador, F.; Martin, A.; López-Menéndez, C.; Moreno-Bueno, G.; Santos, V.; Vázquez-Naharro, A.; Santamaria, P.G.; Morales, S.; Dubus, P.R.; Muinelo-Romay, L.; et al. Lysyl Oxidase–like Protein LOXL2 Promotes Lung Metastasis of Breast Cancer. Cancer Res. 2017, 77, 5846–5859, doi:10.1158/0008-5472.CAN-16-3152.

37. Cuevas, E.P.; Eraso, P.; Mazón, M.J.; Santos, V.; Moreno-Bueno, G.; Cano, A.; Portillo, F. LOXL2 drives epithelial-mesenchymal transition via activation of IRE1-XBP1 signalling pathway. Sci. Rep. 2017, 7, 44988, doi:10.1038/srep44988.

38. Jia, D.; Jolly, M.K.; Tripathi, S.C.; Hollander, P. Den; Huang, B.; Lu, M.; Celiktas, M.; Ramirez-Pena, E.; Ben-Jacob, E.; Onuchic, J.N.; et al. Distinguishing Mechanisms Underlying EMT Tristability. Cancer Converg. 2017, 1, 2, doi:10.1101/098962.

39. Chakraborty, P.; George, J.T.; Tripathi, S.; Levine, H.; Jolly, M.K. Comparative Study of Transcriptomics-Based Scoring Metrics for the Epithelial-Hybrid-Mesenchymal Spectrum. Front. Bioeng. Biotechnol. 2020, 8, doi:10.3389/fbioe.2020.00220.

40. Jones, C.A.R.; Cibula, M.; Feng, J.; Krnacik, E.A.; McIntyre, D.H.; Levine, H.; Sun, B. Micromechanics of cellularized biopolymer networks. Proc. Natl. Acad. Sci. 2015, 112, E5117–E5122, doi:10.1073/pnas.1509663112.

41. Feng, J.; Levine, H.; Mao, X.; Sander, L.M. Nonlinear elasticity of disordered fiber networks. Soft Matter 2016, 12, 1419–1424, doi:10.1039/c5sm01856k.

42. Broedersz, C.P.; Mao, X.; Lubensky, T.C.; Mackintosh, F.C. Criticality and isostaticity in fibre networks. Nat. Phys. 2011, 7, 983–988, doi:10.1038/NPHYS2127.

43. Jansen, K.A.; Licup, A.J.; Sharma, A.; Rens, R.; MacKintosh, F.C.; Koenderink, G.H. The Role of Network Architecture in Collagen Mechanics. Biophys. J. 2018, 114, 2665–2678, doi:10.1016/j.bpj.2018.04.043.

44. Sharma, A.; Licup, A.J.; Jansen, K.A.; Rens, R.; Sheinman, M.; Koenderink, G.H.; Mackintosh, F.C. Strain-controlled criticality governs the nonlinear mechanics of fibre networks. Nat. Phys. 2016, 12, 584–587, doi:10.1038/nphys3628.

45. Rice, A.J.; Cortes, E.; Lachowski, D.; Cheung, B.C.H.; Karim, S.A.; Morton, J.P.; del Río Hernández, A. Matrix stiffness induces epithelial–mesenchymal transition and promotes chemoresistance in pancreatic cancer cells. Oncogenesis 2017, 6, e352–e352, doi:10.1038/oncsis.2017.54.

46. Lu, M.; Jolly, M.K.; Levine, H.; Onuchic, J.N.; Ben-Jacob, E. MicroRNA-based regulation of epithelial-hybrid-mesenchymal fate determination. Proc. Natl. Acad. Sci. 2013, 110, 18144–18149, doi:10.1073/pnas.1318192110.

47. Gregory, P.A.; Bracken, C.P.; Smith, E.; Bert, A.G.; Wright, J.A.; Roslan, S.; Morris, M.; Wyatt, L.; Farshid, G.; Lim, Y.-Y.; et al. An autocrine TGF-β/ZEB/miR-200 signaling network regulates establishment and maintenance of epithelial-mesenchymal transition. Mol. Biol. Cell 2011, 22, 1686–1698, doi:10.1091/mbc.e11-02-0103.

48. Scheel, C.; Eaton, E.N.; Li, S.H.-J.; Chaffer, C.L.; Reinhardt, F.; Kah, K.-J.; Bell, G.; Guo, W.; Rubin, J.; Richardson, A.L.; et al. Paracrine and Autocrine Signals Induce and Maintain Mesenchymal and Stem Cell States in the Breast. Cell 2011, 145, 926–940, doi:10.1016/j.cell.2011.04.029.

49. Katsuno, Y.; Meyer, D.S.; Zhang, Z.; Shokat, K.M.; Akhurst, R.J.; Miyazono, K.; Derynck, R. Chronic TGF-β exposure drives stabilized EMT, tumor stemness, and cancer drug resistance with vulnerability to bitopic mTOR inhibition. Sci. Signal. 2019, 12, eaau8544, doi:10.1126/scisignal.aau8544.

50. Jia, W.; Deshmukh, A.; Mani, S.A.; Jolly, M.K.; Levine, H. A possible role for epigenetic feedback regulation in the dynamics of the epithelial–mesenchymal transition (EMT). Phys. Biol. 2019, 16, 066004, doi:10.1088/1478-3975/ab34df.

51. Ahn, S.G.; Dong, S.M.; Oshima, A.; Kim, W.H.; Lee, H.M.; Lee, S.A.; Kwon, S.; Lee, J.; Lee, J.M.; Jeong, J.; et al. LOXL2 expression is associated with invasiveness and negatively influences survival in breast cancer patients. Breast Cancer Res. Treat. 2013, 141, 89–99, doi:10.1007/s10549-013-2662-3.

52. Du, X.-G.; Zhu, M.-J. Clinical relevance of lysyl oxidase-like 2 and functional mechanisms in glioma. Onco. Targets. Ther. 2018, Volume 11, 2699–2708, doi:10.2147/OTT.S164056.

53. Cao, C.; Lin, S.; Zhi, W.; Lazare, C.; Meng, Y.; Wu, P.; Gao, P.; Wei, J.; Wu, P. LOXL2 Expression Status Is Correlated With Molecular Characterizations of Cervical Carcinoma and Associated With Poor Cancer Survival via Epithelial-Mesenchymal Transition (EMT) Phenotype. Front. Oncol. 2020, 10, doi:10.3389/fonc.2020.00284.

54. Chang, H.-Y.; Tseng, Y.-K.; Chen, Y.-C.; Shu, C.-W.; Lin, M.-I.; Liou, H.-H.; Fu, T.-Y.; Lin, Y.-C.; Ger, L.-P.; Yeh, M.-H.; et al. High snail expression predicts a poor prognosis in breast invasive ductal carcinoma patients with HER2/EGFR-positive subtypes. Surg. Oncol. 2018, 27, 314–320, doi:10.1016/j.suronc.2018.05.002.

55. Drápela, S.; Bouchal, J.; Jolly, M.K.; Culig, Z.; Souček, K. ZEB1: A Critical Regulator of Cell Plasticity, DNA Damage Response, and Therapy Resistance. Front. Mol. Biosci. 2020, 7, doi:10.3389/fmolb.2020.00036.

56. Tripathi, S.; Levine, H.; Jolly, M.K. The Physics of Cellular Decision Making During Epithelial–Mesenchymal Transition. Annu. Rev. Biophys. 2020, 49, 1–18, doi:10.1146/annurev-biophys-121219-081557.

57. Bocci, F.; Gearhart-Serna, L.; Boareto, M.; Ribeiro, M.; Ben-Jacob, E.; Devi, G.R.; Levine, H.; Onuchic, J.N.; Jolly, M.K. Toward understanding cancer stem cell heterogeneity in the tumor microenvironment. Proc. Natl. Acad. Sci. 2019, 116, 148–157, doi:10.1073/pnas.1815345116.

