## Supplemental Information Text for "A theoretical approach to coupling the epithelial-mesenchymal transition (EMT) to extracellular matrix (ECM) stiffness via LOXL2"

### LOXL2 correlation analysis

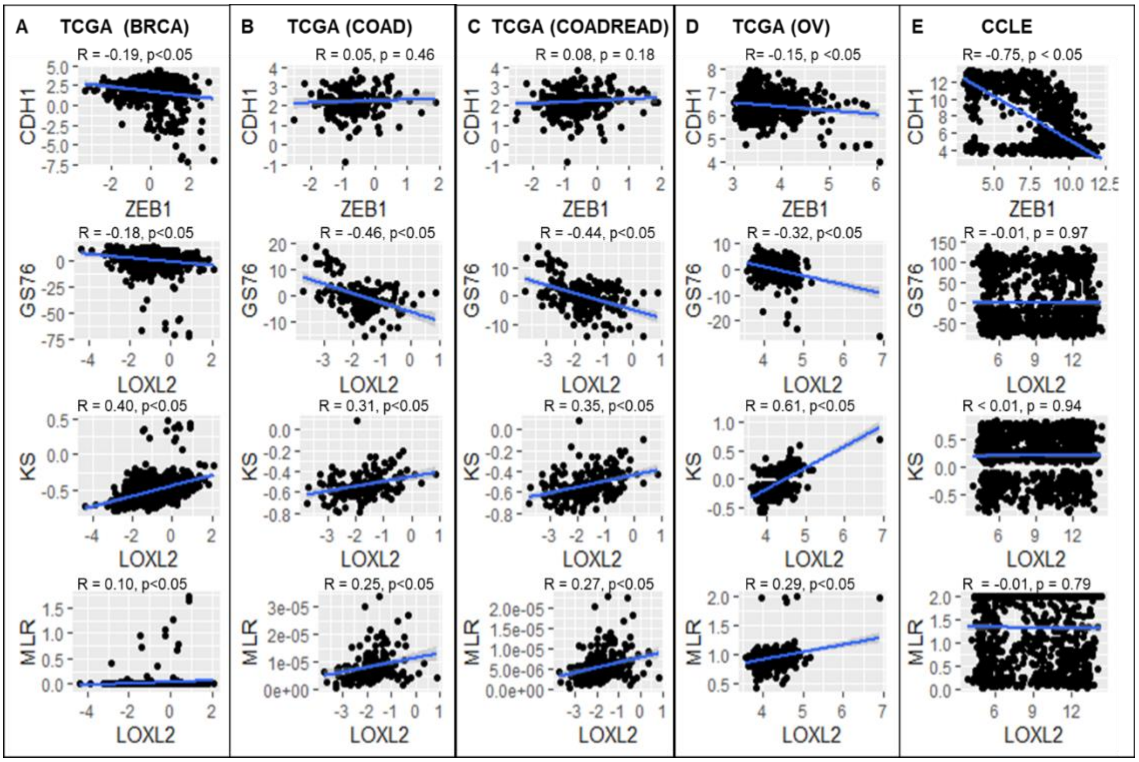

*Figure S1: Correlation between EMT scores using 76GS, KS and MLR methods and LOXL2 gene expression values. Pearson's correlation values (R, p) are been included; the regression line has been highlighted in blue.*

### Mechanical lattice network

In the simulation of bulk shearing, a 60-by-60 triangular lattice with periodic boundary condition is used. Recall that

$$E_{stretch}(i,j) = \frac{1}{2}k\left(l_{ij} - 1\right)^2,$$
$$E_{bend}(i,j,k) = \frac{1}{2}\kappa\left(\theta_{ijk}\right)^2.$$

The stiffness constants $k = 1, \kappa = 1e - 3$.

In the simulation of bead displacement to measure local stiffness, the same values of $k, \kappa$ were used, but we used a 100-by-100 lattice with fixed boundary condition. The fixed boundary here is more sensible because under periodic boundary condition the whole lattice would be dragged along with the single bead with no local deformation. Recall that

$$E_{trap} = \frac{1}{2}k_{trap}\left(d_{bead}^{\parallel} - d_{trap}^{\parallel}\right)^2.$$

Here, $k_{trap} = 0.005,\ d_{trap}^{\parallel} = 1.8$. The bead is of circular shape of radius 1.5. Throughout the network, $p_{bond} = 0.8$. For $p_{phan}$ recall

$$p_{phan} = \frac{p_{phan}^{(0)}}{1 + \left(\frac{c_{LOXL}}{c_{LP}^{(0)}}\right)^2},$$

$$c_{LOXL} = \alpha\rho \text{ if } r < R$$

$$c_{LOXL} = \alpha\rho \frac{R}{r} e^{-(r-R)/d} \text{ if } r > R$$

Here, $p_{phan}^{(0)} = 0.9, c_{LP}^{(0)} = \frac{1}{\sqrt{8}}, \alpha\rho = 1, R = 12.5, d = 3.125.$

In those bead measurements, a smaller area of 85-by-85 central to the whole lattice is measured to avoid edge effects. This is shown in Figure S2.

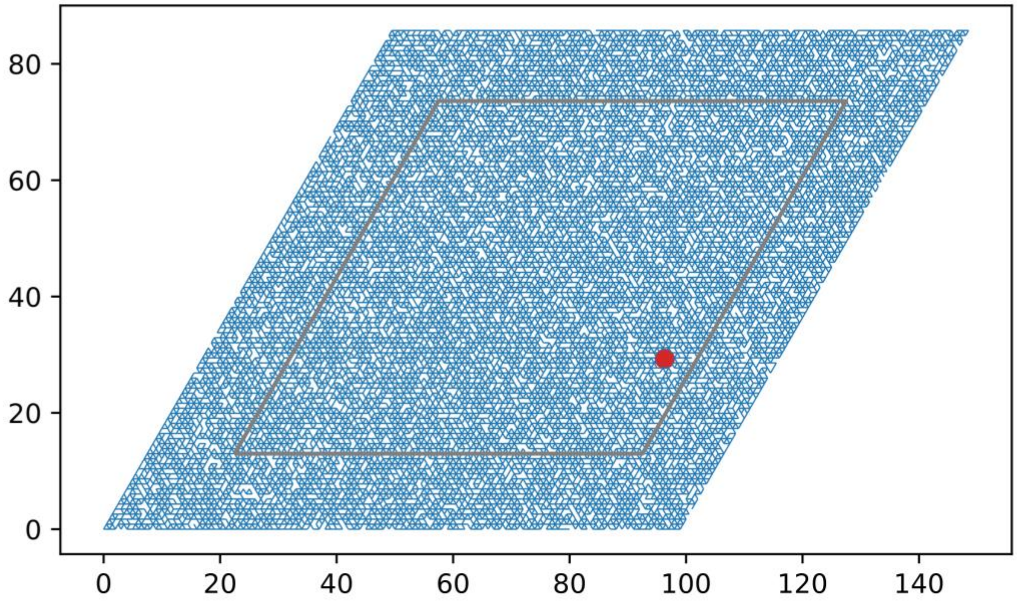

*Figure S2: Bead measurement area of the whole lattice shown in grey box. An 85-by-85 area is measured in the 100-by-100 lattice.*

**Biochemical circuit of EMT**

The full system of ordinary differential equations governing the biochemical network is listed as follows.

$$\dot{\mu}_{200} = g_{\mu_{200}} H^s(Z, \lambda_{z,\mu_{200}}) H^s(S, \lambda_{S,\mu_{200}}) - m_z Y_\mu(\mu_{200}) - k_\mu \mu_{200}$$

$$\dot{m}_z = g_{m_z} H^s(Z, \lambda_{z,m_z}) H^s(S, \lambda_{s,m_z}) - m_z Y_m(\mu_{200}) - k_{m_z} m_z$$

$$\dot{Z} = g_z m_Z L(\mu_{200}) - k_z Z$$

$$\dot{\mu}_{34} = g_{\mu_{34}} H^s(S, \lambda_{s,\mu_{34}}) H^s(Z, \lambda_{z,\mu_{34}}) - m_s Y_\mu(\mu_{34}) - k_{\mu_{34}} \mu_{34}$$

$$\dot{m}_s = g_{m_s} H^s(S, \lambda_{s,m_s}) H^s(I + I_{ECM}, \lambda_{I,m_s}) - m_s Y_m(\mu_{34}) - k_{m_s} m_s$$

$$\dot{S} = g_s m_s L(\mu_{34}) - k_s S$$

$$\dot{c}_{LOXL} = \frac{g_{LOXL}}{1 + \left(\frac{K_{ZL}}{Z}\right)^{n_{ZL}}} - k_{LOXL} c_{LOXL}$$

$$I_{ECM} = \frac{g_{ECM}}{1 + \left(\frac{K_{LE}}{c_{LOXL}}\right)^{n_{LE}}}$$

Here, functions $Y$, $L$ are defined through microRNA-based dynamics, whose precise definition can be found at [1]. $H^S$ is the shifted Hill function defined as $H^S = H^- + \lambda H^+$, where $H^-(B) = \frac{1}{1+\left(\frac{B}{B_0}\right)^{n_B}}$, $H^+ = 1 - H^-$. $\lambda > 1$ corresponds to activation, while $\lambda < 1$ to inhibition.

Table S1 shows the relative parameters in these ODEs.

*Table S1: Parameters in the EMT circuit*

In shifted Hill functions

| Description | Fold change | Value | # of binding sites | Value | Threshold | Value (K molecules) |
|---|---|---|---|---|---|---|
| Inhibition on miR-200 by ZEB | $\lambda_{Z,\mu_{200}}$ | 0.1 | $n_{Z,\mu_{200}}$ | 3 | $Z^0_{\mu_{200}}$ | 220 |
| Inhibition on miR-200 by SNAIL | $\lambda_{S,\mu_{200}}$ | 0.1 | $n_{S,\mu_{200}}$ | 2 | $S^0_{\mu_{200}}$ | 180 |
| Self-activation of ZEB | $\lambda_{Z,m_z}$ | 7.5 | $n_{Z,m_z}$ | 2 | $Z^0_{m_z}$ | 25 |

| | | | | | | |
|---|---|---|---|---|---|---|
| Activation on ZEB by SNAIL | $\lambda_{S,m_z}$ | 10.0 | $n_{S,m_z}$ | 2 | $S^0_{m_z}$ | 180 |
| Inhibition on miR-34 by SNAIL | $\lambda_{S,\mu_{34}}$ | 0.1 | $n_{S,\mu_{34}}$ | 1 | $S^0_{\mu_{34}}$ | 300 |
| Inhibition on miR-34 by ZEB | $\lambda_{Z,\mu_{34}}$ | 0.2 | $n_{Z,\mu_{34}}$ | 2 | $Z^0_{\mu_{34}}$ | 600 |
| Self-inhibition of SNAIL | $\lambda_{S,m_s}$ | 0.1 | $n_{S,m_s}$ | 1 | $S^0_{m_s}$ | 200 |
| Activation on SNAIL by external signal I | $\lambda_{I,m_s}$ | 10 | $n_{I,m_s}$ | 2 | $I^0_{m_s}$ | 50 |

In function $Y$ and $L$

| n (# of miRNA binding sites) | 0 | 1 | 2 | 3 | 4 | 5 | 6 |
|---|---|---|---|---|---|---|---|
| $l_i$(hour$^{-1}$) | 1 | 0.6 | 0.3 | 0.1 | 0.05 | 0.05 | 0.05 |
| $\gamma_{mi}$(hour$^{-1}$) | 0 | 0.04 | 0.2 | 1 | 1 | 1 | 1 |
| $\gamma_{\mu i}$(hour$^{-1}$) | 0 | 0.005 | 0.05 | 0.5 | 0.5 | 0.5 | 0.5 |
| $n_{\mu_{200}}$ | 6 | $n_{\mu_{34}}$ | | | | | 2 |
| $\mu^0_{200}$ | 10K | $\mu^0_{34}$ | | | | | 10K |

Other rate constants

| Synthesis rate | Value (molecules/hour) | Degradation rate | Value (hour$^{-1}$) | Translation rate | Value (hour$^{-1}$) |
|---|---|---|---|---|---|
| $g_{\mu_{200}}$ | 2.1K | $k_{\mu_{200}}$ | 0.05 | $g_z$ | 0.1K |
| $g_{m_z}$ | 11 | $k_{m_z}$ | 0.5 | $g_s$ | 0.1K |

| | | | |
|---|---|---|---|
| $g_{\mu_{34}}$ | 1.35K | $k_z$ | 0.1 |
| $g_{m_s}$ | 90 | $k_{\mu_{34}}$ | 0.05 |
| | | $k_{m_s}$ | 0.5 |
| | | $k_s$ | 0.125 |

In LOXL2 and ECM pathways

| Symbol | Value | Unit |
|---|---|---|
| $g_{LOXL}$ | 0.1 | K molecules / hour |
| $k_{LOXL}$ | 0.1 | hour$^{-1}$ |
| $K_{ZL}$ | 250 | K molecules |
| $n_{ZL}$ | 1 | 1 |
| $g_{ECM}$ | 80 | K molecules |
| $K_{LE}$ | varied | K molecules |
| $n_{LE}$ | 4 | 1 |

**Bifurcation diagrams**

In main text Figure 4 we showed the family of bifurcation diagrams as the parameter $K_{LE}$ is varied between closed interval $[0, 1]$ with step increase of $0.1$. If $K_{LE} = 0$, only the mesenchymal steady state exists (Figure S3). If $K_{LE} = 1$, the bifurcation is barely affected by the new LOXL2 edge (Figure S5). In some intermediate value such as $K_{LE} = 0.5$, the mesenchymal state is stabilized and can even bootstrap itself under zero external driving signal (Figure S4). See main text for discussion.

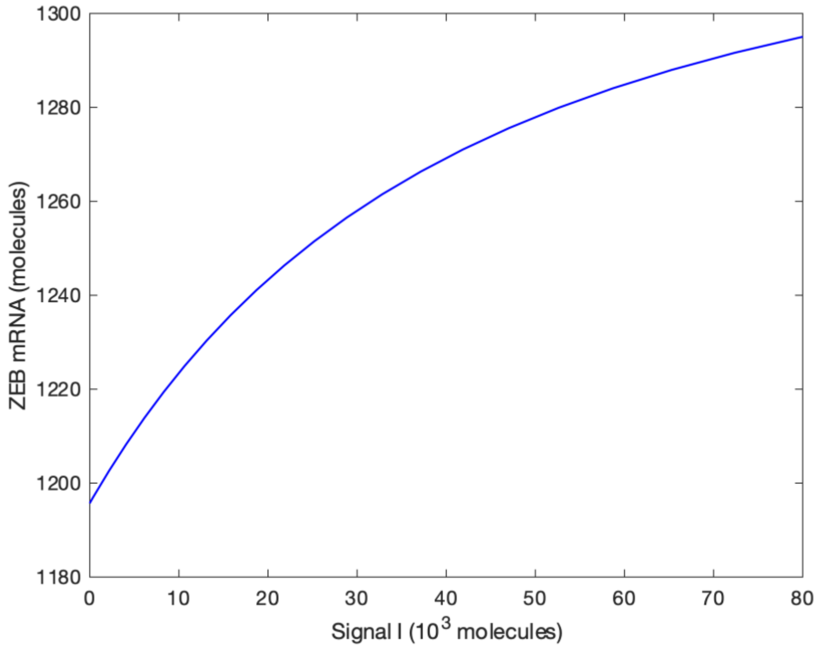

*Figure S3: Bifurcation diagram when* $K_{LE} = 0$.

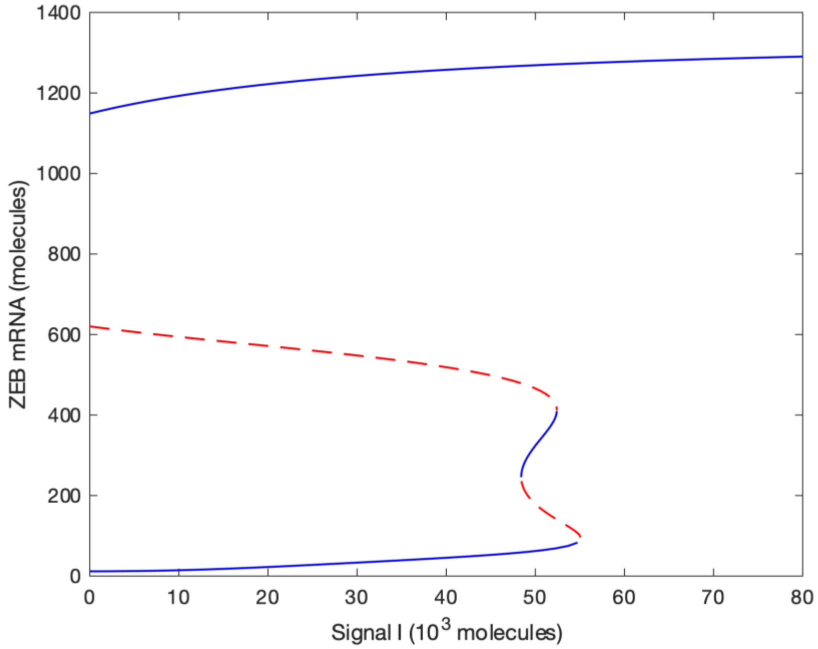

*Figure S4: Bifurcation diagram when* $K_{LE} = 0.5$.

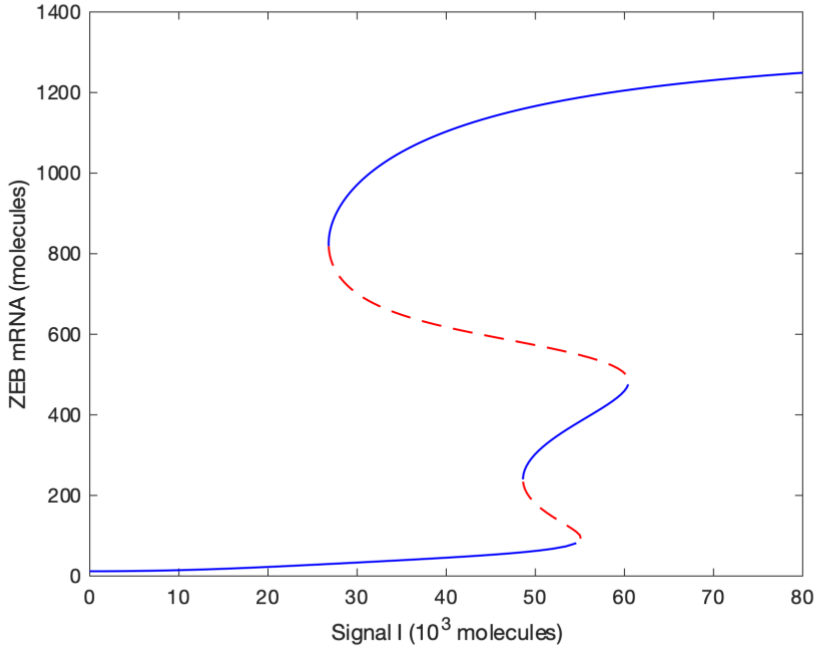

*Figure S5: Bifurcation diagram when $K_{LE} = 1$.*

1. Lu, M.; Jolly, M.K.; Levine, H.; Onuchic, J.N.; Ben-Jacob, E. MicroRNA-based regulation of epithelial-hybrid-mesenchymal fate determination. *Proc. Natl. Acad. Sci.* **2013**, *110*, 18144–18149, doi:10.1073/pnas.1318192110.
